# Trichome formation in *Nicotiana benthamiana* is induced by *Agrobacterium*

**DOI:** 10.64898/2025.12.02.691950

**Authors:** Jian Chen, Phil Hands, Manish Patel, Leilei Yang, Chengcheng Zhong, Neil Smith, Ming Luo, Michael Ayliffe

## Abstract

Plant trichomes are specialised epidermal appendages found on various organs of many plant species. They play important roles in plant protection against pests, pathogens, herbivores, resistance to abiotic stress, light perception, mechanosensitivity etc. They are of direct agricultural importance, providing fibre (e.g. cotton) and producing secondary metabolites essential for agriculture, medicine, and industry. Trichomes can be relatively sparsely distributed, making their biological investigation and commercial exploitation more challenging. Here we show that infiltration of *Nicotiana benthamiana* leaf tissue with certain nopaline-type *Agrobacterium tumefacians* strains induces extremely dense, localised formation of glandular trichomes 15 days post infiltration (dpi). This *Agrobacterium*-strain specific effect was shown to be due to the presence of a *trans-zeatin synthase* (*tzs*) gene located on some Ti plasmids, which results in the bacterial production of the cytokinin trans-zeatin. This simple procedure enables dramatically increased trichome density to be compared biologically with adjacent isogenic plant tissues on the same leaf.

## Results and Discussion

*N. benthamiana* leaf tissue (abaxial side) was infiltrated with *A. tumefacians* strain GV3101 PMP90, which contains disarmed Ti plasmid pTiC58PMP90 (GenBank KY000034). Three leaves on 4 plants were infiltrated with 4 sites per leaf. Fourteen dpi abundant adaxial trichome formation was visible at infiltration sites that was absent in adjacent uninfiltrated tissue (Figure 1A, panel 1; Figure S1). Infiltrations were repeated and scanning electron microscopy (SEM) showed the vast majority of trichomes at infiltration sites were multicellular, capitate glandular trichomes, previously described on abaxial and adaxial leaf surfaces of *N. benthamiana* (Slocombe et al. 2008) (Figure 1A, panels 2 and 3). 6 dpi emergent trichomes were observed on epidermal pavement cells with multiple trichomes on some cells that matured 12-15 dpi (Figure 1A, panels 4 and 5). An increase in abaxial trichome density was also observed at infiltration sites, although not as dense as on the adaxial surface (Figure S2).

**Figure 1:**
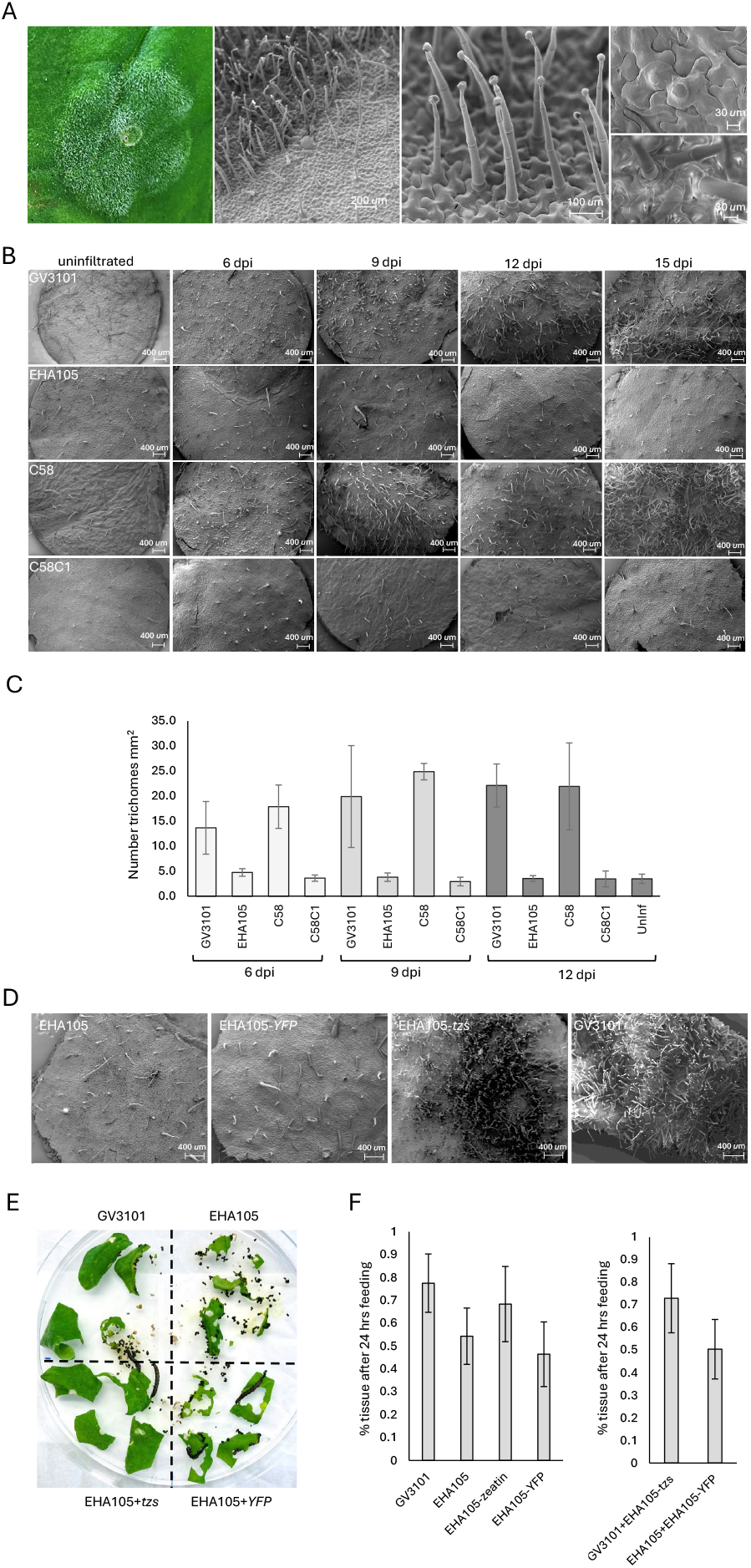
a) From left to right: A GV3101 infiltration site 15 dpi on a *N. benthamiana* leaf showing abundant visible trichome formation; SEM image of a GV3101 infiltration site boundary zone with high trichome density in the infiltration site and reduced trichome density in adjacent uninfiltrated tissue; multicellular, capitate glandular trichomes produced after GV3101 infiltration; (upper) two emergent trichomes produced from a single epidermal pavement cell; multiple mature trichomes formed from epidermal pavements cells 15 dpi. b) SEM time course of *N. benthaniana* leaves following infiltration with strains GV3101, EHA105, C58 and C58C1. c) Average trichome densities at *Agrobacterium* infiltration sites. Standard deviations shown. d) SEM images 15 dpi with strains EHA101, EHA101-*YFP*, EHA105-*tzs* and GV3101. e) *Helicoverpa armigera* feeding after 24 hours on *N. benthamiana* leaf segments previously infiltrated (10 dpi) with either GV3101, EHA105, EHA105-*tzs* or EHA105-*YFP*. f) % of infiltrated leaf tissue remaining 24 hours after *H. armigera* feeding. *Agrobacterium* strains are indicated on the X axis. The graph at right shows combined data from trichome producing infiltrations (GV3101 and EHA105-*tzs*) and non-trichome inducing infiltrations (EHA015 and EHA105-*YFP*).

Experiments were repeated using strain GV3101 pMP90 and strains EHA105, C58 and C58C1. These latter *Agrobacterium* carry disarmed Ti plasmid pTiBo542Δ, wild type Ti plasmid pTiC58 (GenBank NZ_KY000040) and no Ti plasmid, respectively. After leaf infiltration a 6, 9, 12 and 15 dpi SEM time course was undertaken. Six dpi numerous small emergent trichomes were observed at GV3101 pMP90 and C58 infiltration sites in juxtaposition to larger pre-existing mature trichomes, that were also present on uninfiltrated control tissue (Figure 1B). Over time more emergent trichomes were observed that continued to grow forming a dense patch 15 dpi (Figure 1B). In contrast, EHA105 and C58C1 infiltration did not induce trichome formation with only sparse pre-existing mature trichomes observed (Figure 1B). Trichomes density was 6 times greater on GV3101 pMP90 and C58 infiltrated sites 12 dpi compared with other sites (Figure 1C).

Trichome induction by strain C58 but not C58C1, which is a C58 derivative cured of the wildtype Ti plasmid pTiC58, suggests Ti plasmid encoded gene(s) cause the induction. GV3101 and EHA105 are both derivatives of strain C58, hence having similar genomic backgrounds, but each carries a different disarmed Ti plasmid, further suggesting Ti plasmid gene(s) induce trichome formation. Wildtype Ti plasmids pTiC58 and pTiBo54 (GenBank NC_010929), modified to produce disarmed Ti plasmids in GV3101 and EHA105, are from different lineages (Weisberg et al. 2020).

Cytokinins can induce trichome formation when applied to plant leaf tissue (Maes and Goossens, 2010). *Agrobacterium* produces cytokinin in two ways. Firstly, by transfer of the T-DNA encoded *isopentenyl transferase* (*ipt*) gene into the plant host (Barry et al. 1984). However, *ipt* is absent from pTiC58PMP90 due to disarmament, precluding this gene from being causative. The second cytokinin source is direct bacterial production of trans-zeatin by a *trans-zeatin synthase* (*tzs*) gene located in the Ti plasmid *vir* region (Beaty et al. 1986). pTiC58 and pTiC58PMP90 encode identical 732 bp *tzs* ORFs at nucleotides 179993-179262 and 127756-127025, respectively. No *tzs* gene is encoded by pTiBo542, although the pTiC58 tzs protein and pTiBo542 ipt protein show 53% identity (Beaty et al. 1986). GV3101 PMP90 was previously shown to induce shoot regeneration in tissue culture in the absence of cytokinin whereas EHA105 could not (Han et al. 2013).

A 4.3 kb fragment (Figure S3) encoding *tzs* (Erickson et al. 2014) was PCR amplified from pTiC58PMP90 and inserted into a pCSIRO binary vector with T-DNA border sequences removed. This construct replicates in *Agrobacterium* but is incapable of T-DNA transfer. The pCSIRO-*tzs* plasmid was introduced into EHA105 to create EHA105-*tzs* which induced abundant trichome development in *N. benthamiana* upon infiltration (Figure 1D). Therefore, the *tzs* gene is causative for the observed trichome production. To further confirm the role of cytokinin in the observed phenotype 10-100 ug/ml of 6-benzylaminopurine was infiltrated into *N. benthamiana* leaves and increased trichome formation was observed at higher concentrations, although less dense than by GV3101 (Figure S4).

The induction of trichomes by *tzs* expressing *Agrobacterium* enables trichome effects to be compared between isogenic tissues on the same leaf and at the same developmental stage. Insect feeding experiments were undertaken on *N. bethamiana* leaf segments infiltrated with either strain GV3101, EHA105, EHA105-*tzs* or EHA105 carrying a *35S-YFP* encoding binary T-DNA vector (EHA105-*YFP*). Infiltrated leaf regions were excised 10 dpi and placed on MS media. Each plate quadrant contained leaf segments infiltrated with the same bacterial strain (Figure 1E). Five 2nd-instar *Helicoverpa armigera* (cotton boll worm) larvae were placed on each of five biological replicate plates and the % leaf tissue remaining in each quadrant quantified after 24 hours (36 hours in one experiment due to delayed larval feeding) (Figure S5). Tissue infiltrated with either strain GV3101 or EHA105-*tzs* showed a reduced insect feeding trend when compared with EHA105 or EHA105-*YFP*, although values were not significantly different (Figure 1F). When data from trichome inducing and non-inducing strains was pooled a highly significant difference in insect eating preference was apparent (P=0.002, students T-test) (Figure 1F).

This insect feeding experiment is exemplar of studies that can be undertaken using isogenic tissues with extreme differences in trichome density. This trichome assay will be beneficial in dissecting the cellular developmental and biological role of trichomes in addition to trichome biofactory development for valuable plant secondary metabolite production.

## Acknowledgements

We thank Dr Ian Greaves for supplying *Agrobacterium* strains C58 and C58C1. This work was funded by CSIRO.

## Author contributions

J.C., P.H., M.P., N.S., M.L and M.A participated in the conception and design of the research. J.C., P.H., M.P., N.S., M.L. undertook experiments. C.Z. and L.Y. were involved in insect feeding experiments. All authors contributed to writing the manuscript. All authors agreed to the submitted version of the manuscript.

## Supplementary data

**Figure S1:** Examples of *N. benthamina* trichome production after infiltration with *Agrobacterium* strain GV3101.

**Figure S2:** Comparison of trichome densities on abaxial and adaxial leaf surfaces after GV3101 infiltration.

**Figure S3:** Sequence of the 4.3 kb fragment encoding the *tzs* gene introduced into *Agrobacterium* strain EHA105.

**Figure S4:** 6-benzylaminopurine induces trichome formation in *N. benthamiana*.

**Figure S5:** Replicate plates used for *Helicoverpa armigera* feeding trials.

## Methods

### Plant propagation

*N. benthamiana* plants were grown under controlled environment conditions (23° with 16⍰h light:⍰8⍰h dark photoperiod) for 3-4⍰weeks in a 3:1 Scotts Premium Mix and Debco soil with 4 g/L Osmocote. Insects were controlled by hanging sticky insect traps in the plant growth environment.

### Agrobacterium infiltration assays

Cultures of *Agrobacterium* strains GV3101 pMP90, EHA105, C58 and C58C1 were grown by inoculating single colonies in Luria-Bertani liquid media (10g tryptone, 5g yeast extract and 10g NaCl/L, pH 7.0) containing appropriate antibiotic selection and grown at 28°C with vigorous shaking for 24⍰h. Cultures were then centrifuged at 2500 g for 5 minutes and resuspended in infiltration buffer (10⍰mm MES pH 5.6, 10⍰mm MgCl_2_ and 150⍰μm acetosyringone) at OD_600 nm_ of 0.8. *N. benthamiana* leaves were then syringe infiltrated with *Agrobacterium* solutions using a 1 ml syringe. Generally, 4-6 infiltrations were undertaken on a single leaf and replicate infiltrations were undertaken on multiple leaves from independent plants.

### Production of plasmid pCSIRO-tzs

A 4.3 kb fragment encoding the *tzs* gene (Figure S2) was DNA synthesised (Epoch Life Sciences, USA) and then PCR amplified with a primer pair (primers CCCTTTTAAATATCCGATTATTCTAATAAACGCTCTTT-TCTCATGGCTCATGGCTCAGGCAGCTTCGCA and GATTTTGTGCCGAGCTGCCGGTCGGGGAGCTGTTGGCTG-GCTCTGATAGAGGAGACCAGAGTAACTTGG) containing homology to the backbone sequence of binary vector pCSIRO. pCSIRO was then PCR amplified with primers GAGAAAAGAGCGTTTATTAGAATAATCGG and AGCCAGCCAACAGCTCCCCG which amplify the vector backbone without the T-DNA region or LB and RB sequences. The *tzs* gene was inserted into the pCSIRO backbone by Gibson assembly. This plasmid, *pCSIRO-tzs*, was transformed into *Agrobacterium* strain EHA105 and transformants selected by growth on LB agar containing 25 ug/ml rifampicin and 50 ug/ml kanamycin.

### Scanning electron microscopy

8mm leaf discs were taken from *Agrobacterium* infiltrated areas, cleared overnight in 100% EtOH and transferred to fresh EtOH. Discs were critical point dried in an Autosamdri-815 automatic critical point drier (Tousimis Research Corporation, Rockville USA) and mounted onto aluminium stubs with double-sided adhesive carbon tabs (ProSciTech, Townsville City, QLD, Australia). Samples were gold coated using a MiniQS sputter coater (Quorum Technologies Ltd., EastSussex, UK) for 30 seconds under an argon environment and imaged using a Zeiss EVO LS 15 scanning electron microscope (Carl Zeiss Microscopy GmbH, Jena, Germany) under high vacuum using a secondary electron detector, 2 kV accelerating voltage, spot size 300. For determination of trichome densities, trichomes were counted on SEM images of 4 *Agrobacterium* infiltration sites per strain after 6, 9 and 12 dpi. Trichomes could not be counted at 15 dpi due to the high trichome densities at GV3101 PMP90 and C58 infiltration sites. The 12 dpi calculations are likely an underestimate of final trichome density as more trichome growth was apparent at 15 dpi.

### Propagation of Helicoverpa armigera and feeding assays

*N. benthamiana* was infiltrated with *Agrobacterium* strains of choice and 10 dpi infiltrated leaf tissue excised with a razor blade. Tissues were placed onto 1% agarose (Biobasic, Canada) plates with each quadrant containing tissue infiltrated with the same *Agrobacterium* strain. Plates were then photographed and five *H. armigera* larva at the 2nd instar stage were then added to each plate. A laboratory colony of *H. armigera conferta*, maintained since the mid-1980s, was used for infiltrated leaf feeding trials. Insects were reared at 25 ± 1 °C, 50 ± 10% relative humidity, under a 14 h light: 10 h dark photoperiod to simulate natural conditions. Newly hatched neonates were fed an artificial diet until the second instar stage before being transferred to plates containing *Agrobacterium* infiltrated leaf segments. Plates were then re-photographed after 12, 18, 24 and 36 hours of insect feeding.

The extent of leaf predation was quantified via image analysis using a FIJI image processing package (version 1.53c. Schindelin et al., 2012). Briefly, the region containing each quadrant was manually isolated and a consistent colour threshold applied to isolate only green coloured pixels. For each quadrant region the leaf area, corresponding to green pixels, was quantified and normalised to the area of the entire plate. For each timepoint values were normalised against the area of leaf at the zero time point to determine the % of green tissue remaining.

Five replicate plates were assayed and data from the zero time point and 24 hr time point compared, with the exception of one plate image which was selected after 36 hrs due to slower insect feeding (Figure S5). The % of green tissue remaining was averaged for tissue infiltrated with the same *Agrobacterium* strain and data compared by ANOVA with posthoc Tukey (https://astatsa.com/OneWay_Anova_with_TukeyHSD).

## Supplemental Figures

**Figure S1:**
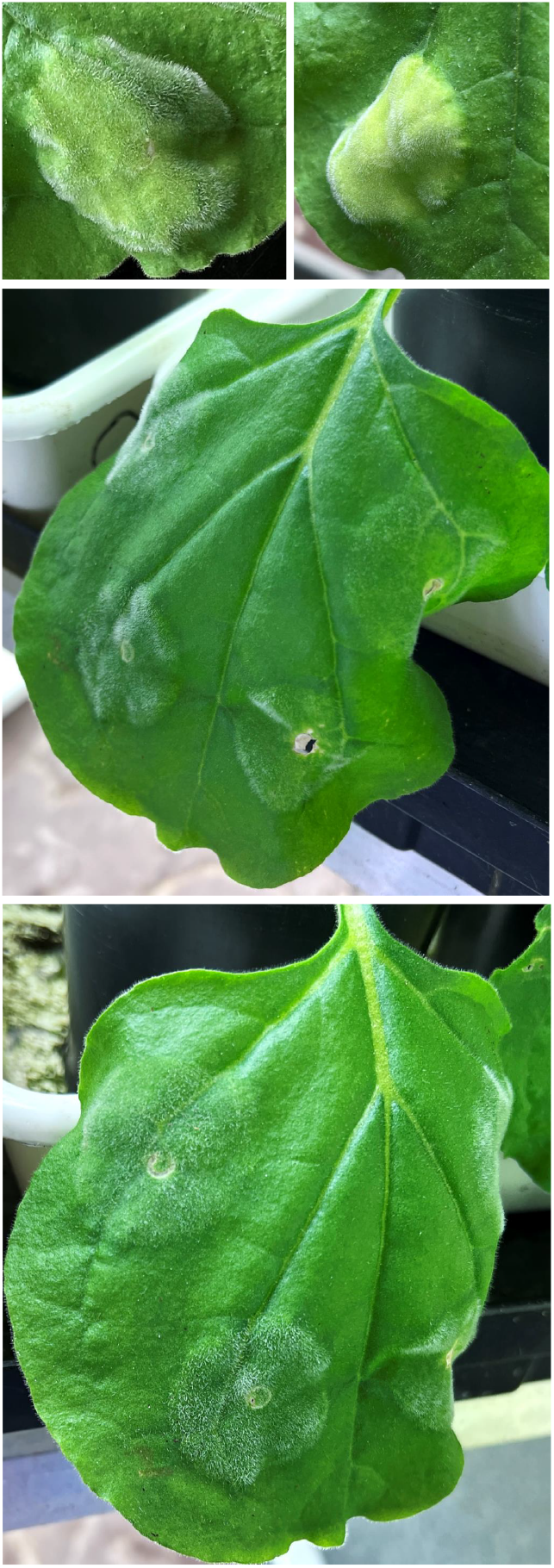
Examples of *N. benthamina* trichome production after infiltration with *Agrobacterium* strain GV3101.

**Figure S2:**
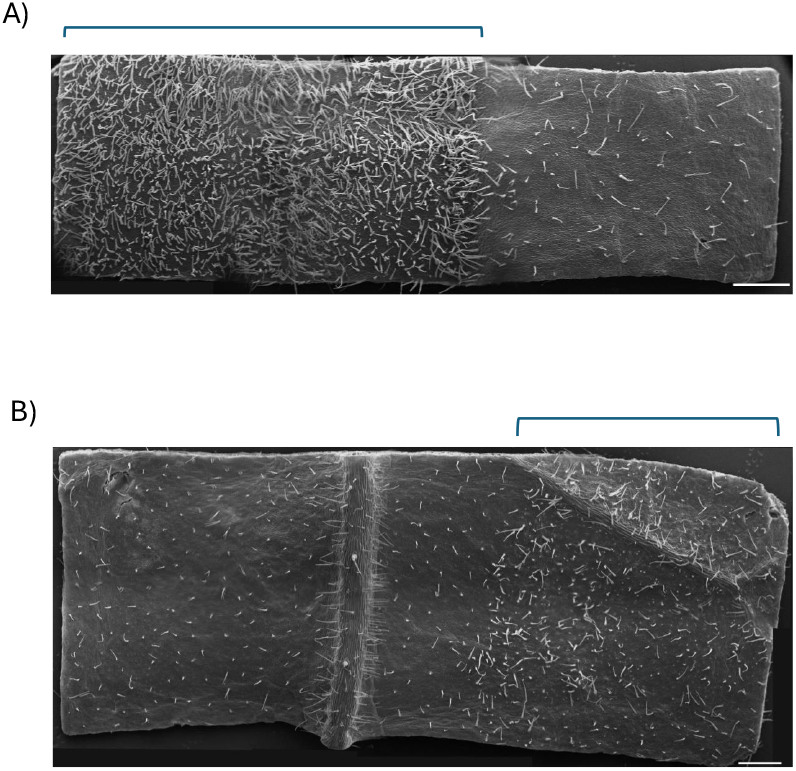
Comparison of trichome densities on adaxial and abaxial leaf surfaces following infiltration with *Agrobacterium* strain GV3101. Panel A shows the adaxial leaf surface while panel B shows the abaxial surface, infiltration zones are marked with brackets and scale bars show 1 mm. Images were assembled from multiple SEM images stitched using the ImageJ plugin, MosaicJ (Thevenaz & Unser, 2007).

**Figure S3:**
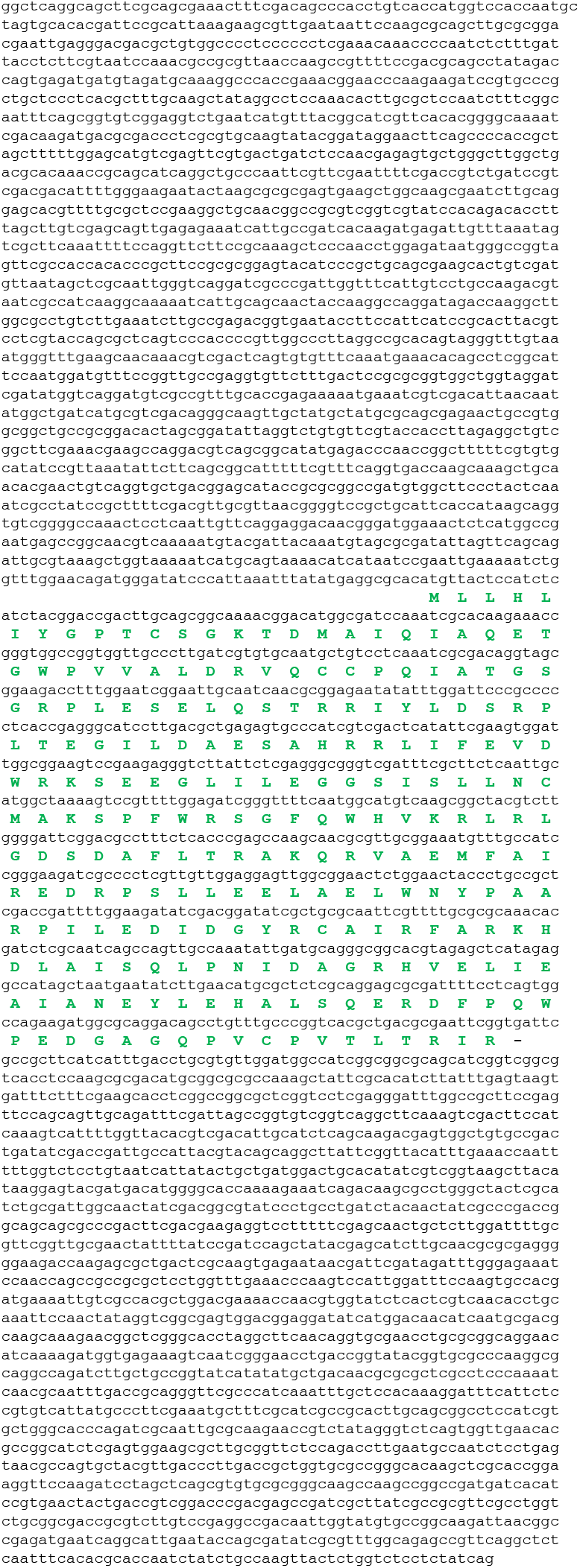
Sequence of the 4.3 kb fragment encoding the *tzs* gene introduced into *Agrobacterium* strain EHA105. The amino acid sequence of the tzs protein is shown in green.

**Figure S4:**
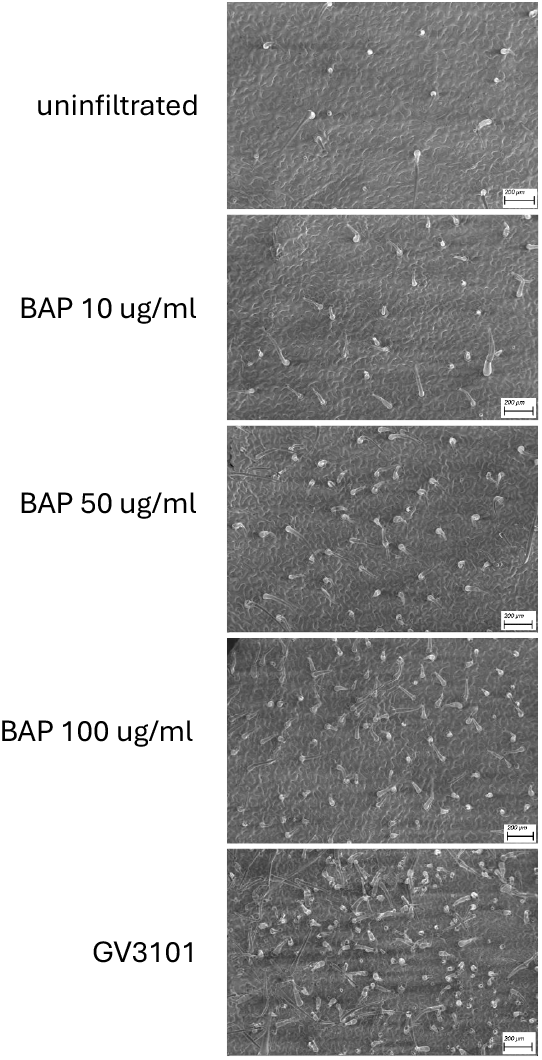
6-benzylaminopurine induces trichome formation in *N. benthamiana*. Leaves were infiltrated with 6-benzylaminopurine concentrations indicated or *Agrobacterium* strain GV3101 and SEM images taken 25 dpi.

**Figure S5:**
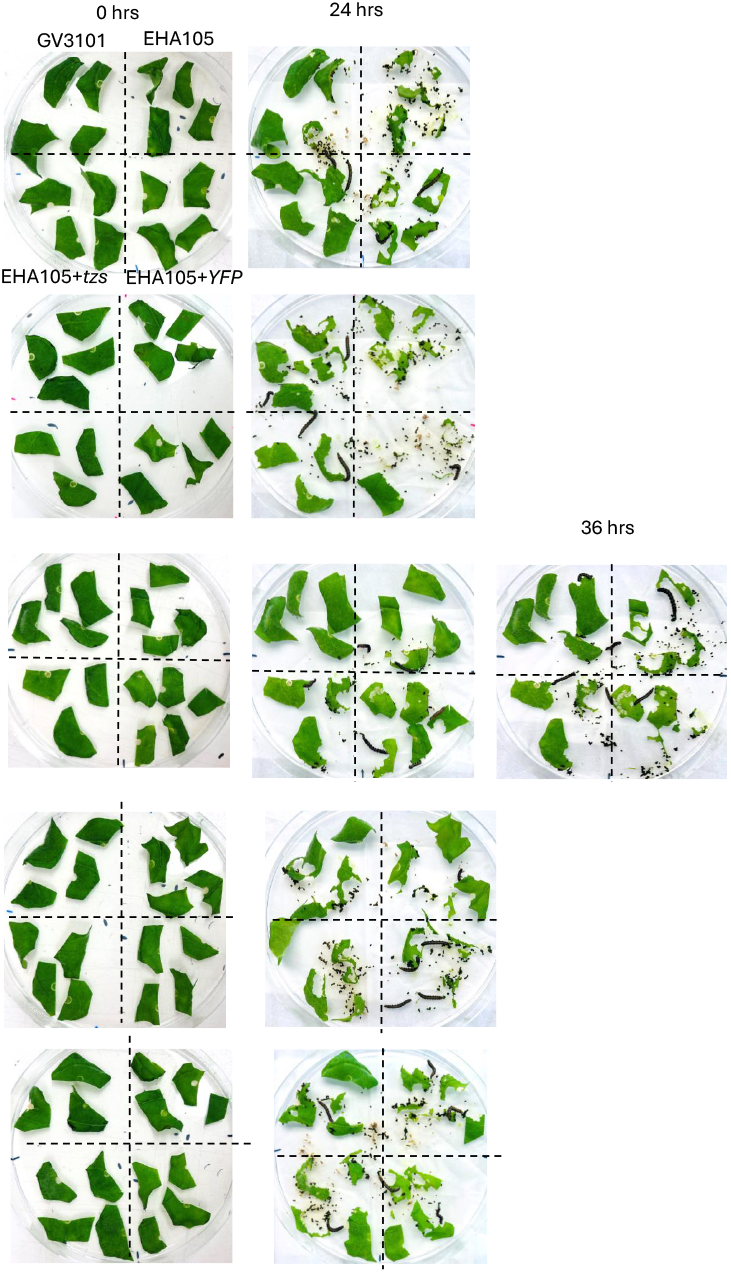
Replicate plates used for *Helicoverpa armigera* feeding trials. Plates at left were prior to larvae addition while plates at right show remaining leaf tissue after 24 hours feeding of the same plate. The third experiment from the top used a 36-hour feeding plate for tissue quantification rather than the 24-hour plate due to delayed insect feeding. Each quadrant contains *N. benthamima* leaf tissue infiltrated with either *Agrobacterium* strain GV3101, EHA105, EHA105+*tzs* or EHA105+*YFP*.

